# Head movement allocation in freely-moving macaques learning object-context sequences

**DOI:** 10.1101/2023.12.11.571113

**Authors:** K. F. Rahman, S. Abbaspoor, W. Zinke, H. Tamaki, K.L. Hoffman

**Author notes:** 2301 Vanderbilt Place, Nashville, TN 37235. These authors contributed equally (K. F. Rahman, S. Abbaspoor). Senior author (K.L. Hoffman).

## Abstract

Flexible learning and long-term memory are hallmark features of primate cognition that are shaped by interactions with changing environments. A comprehensive understanding of these brain-body-environment interactions requires minimal constraints on behavior, and environments rich with conditional cues that encourage diverse behavioral dynamics. In this study, we introduce a multi-level enclosure (‘Treehouse’) that allows for independent control of spatially distributed touchscreens for studying cognition in freely moving macaques. To test flexible learning, two monkeys completed trials consisting of a regular sequence of conditional object choices (‘targets’) across four distinct spatiotemporal contexts (‘screens’). Both monkeys acquired numerous unique, conditional object–context sequences without detriment to new learning as memoranda accumulated. They also successfully recalled their correct choices after weeks without exposure. To assess how monkeys allocate movement during learning and memory-guided decision making, we tracked their head kinematics and used recurrent neural networks to decode different aspects of their engagement with the environment during the choice epoch of the task. One such model predicted the position of the object about to be touched, including on error trials, tracking overt gaze towards objects. Separate models decoded sequence corner, the ordinal step, and the individual screen stations the monkeys occupied, despite identical stimulus layouts and behavioral demands at each station. Head movements also showed efficiencies that the task did not require: under conditions of uncertainty, monkeys were biased in their initial search locations to the targets that were closest to reach, and across days they made smoother head movements as training progressed, with no change in accuracy. These results demonstrate the potential of immersive 3-D environments, in which body kinematics can shed light on animals’ goals, and reveal that situated learning in primates can be rapid, capacious, durable, and flexible.

## Introduction

Memory is organized in the service of adapting behavior to available resources. In macaques, this includes sequentially foraging across distributed locations and flexibly modifying strategies in accordance with the hierarchical structures of their surroundings. This, in turn, requires the development, retention, and generalization of conditional associations across days and weeks, in the face of continual learning. Brain regions playing central roles in this process include the hippocampus and associated neocortex, identified in lesion studies for their contributions to object recognition memory, associative learning and object-place binding^1–9^. Electrophysiological recordings in these areas revealed neural correlates of associative-relational learning abilities^10–17^, though such experiments are typically conducted under constrained testing conditions that lack sensory immersion and restrict natural exploratory movements. These very attributes contribute to performance in foundational lesion studies of hippocampal function in macaques^5,6^ and they are known to benefit associative memory in humans under immersive environments^18–20^. Consequently, under conventional experimental testing designs for macaques, the full conditional structure of primate memory, including its capacity across accumulating memoranda, its resistance to proactive interference, its persistence over weeks, and its dependence on immersive, spatiotemporal context, may have been under-estimated at the scale and complexity that natural behaviors demand.

Free movement also opens an underexploited window onto cognition whereby the body’s own kinematic record may contribute to memory, or to foraging policies that take into account the energetic and cognitive demands that arise under free-moving conditions. For example, kinematics can reveal latent cognitive factors that choice accuracy alone cannot ^21,22^. Even under restricted sampling, where eye movements are the available exploratory movement, saccades and scanpath geometry expose the sequential sampling of alternatives and dynamics of commitment during decisions ^23–25,84^. Furthermore, movement vigor declines as internal models of a task mature, reflecting a shift toward energetically efficient movements with experience ^26–29^ providing a means to evaluate metalearning and strategic refinement ^30^. By increasing freedom of movement, vigor becomes a more distributed property of animal behavior, potentially requiring attention to trackable, quantifiable effectors, and analytic approaches to disentangle movement properties that vary in their energetic efficiency, task-necessity, and policy relevance.

To understand these attributes of learning in natural settings, we constructed a multi-level enclosure (the ‘Treehouse’) equipped with spatially distributed touchscreens. Freely moving macaques were presented sequential objects-in-context to be continually learned as their head kinematics were tracked via an inertial measurement unit.

The enclosure and the task it enabled allow us to pose three general questions. First, can macaques retain spatiotemporally conditional associative memories while continuing to acquire more memoranda? Second, for the degrees of freedom the task doesn’t constrain or demand, are their movements organized according to the task or environment? Third, does the animal economize movement?

## RESULTS

### Learning sequentially presented item-context associations

The testing apparatus consisted of a custom-made enclosure (the ‘Treehouse’, Figure 1A, B) equipped with modular panels organized into upper and lower levels and 8 touchscreens (See Methods for details). Trials consisted of a 4-step sequence, with each step containing a four-object array presented in a predetermined, repeated order across the 4 screens of a corner (Figure 1D). Exactly one object was deemed the ‘Target’ for that step, i.e., screen context, thus it was a distractor for the remaining steps in the sequence (Figure 1C, D). The corner to which a set was assigned was fixed and counterbalanced across sets.

**Figure 1.**
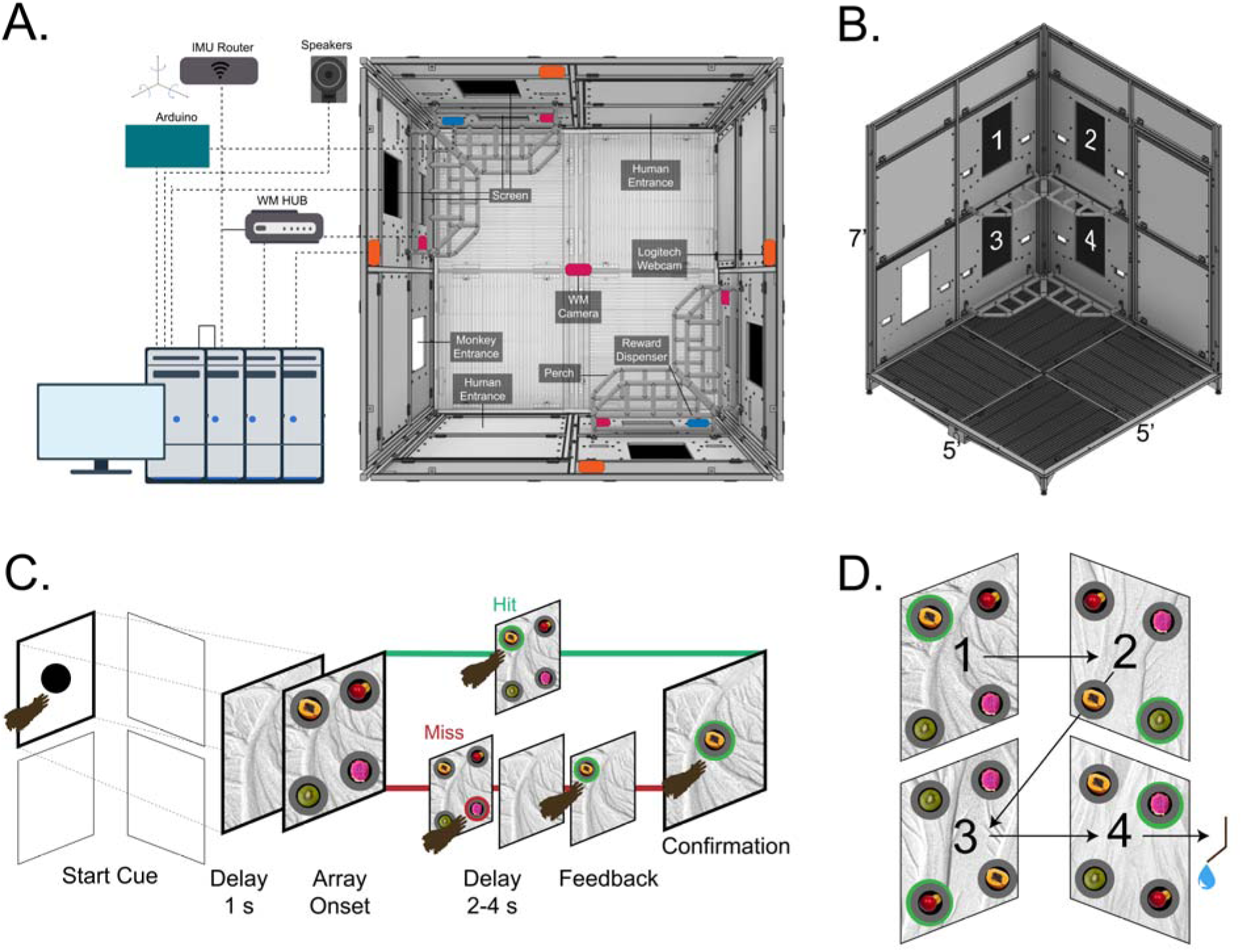
Sequential object-in-context association task in the 3-D ‘Treehouse’ enclosure. (A, B) renderings of the enclosure, including a schematic of peripheral devices that enabled timed stimulus delivery and behavioral measurements. (A) Overhead view of the task environment. Black squares depict touchscreens, white square shows the monkey entrance, blue rectangles show the locations of reward dispensers, and pink rectangles show the camera positions. (B) One corner of the enclosure, revealing the four stations comprising one trial (stations are numbered in presentation order). The opposite corner has been hidden for visibility. (C)Trial sequence depicted for the first screen in the sequence. For illustration only: Bold black = active screen; green ring = correct target; red ring = incorrect distractor. Following completion of one screen’s association, the trial continues to the next screen in the sequence. (D) Object array presentations in an example trial. Green rings indicate the associated targets of screens. The position of the 4 objects in the 2 x 2 array is randomized across screens and trials, and the background image was simplified for illustration. The 4-screen sequence is maintained throughout the study.

The two monkeys completed 14 and 22 sets of 4-item sequences for over 50 and 80 unique item-context memoranda. Completion of each trial required a series of decisions and extensive movement in the enclosure prior to reward. This freedom of movement provided a window into how the body engages with the task; specifically, we examined how head kinematics serves as a proxy for overt attention and how movements signal the location the animal selects, whether it also reflects where the animal is in the enclosure and in the sequence, and the cost of producing that movement. Trial durations (the elapsed time from touch on the first screen cue to when animals completed performance on the 4^th^ screen, triggering reward) were, on average, 40.5 (s.d.11.0) and 29.0 s (s.d. 5.5) for the two monkeys, W and F, respectively (N: 1408 and 2157 trials).

We predicted that (H1) conditional object-in-context memory would be acquired and maintained over weeks’ delay in the face of nested interference. Figures 2A and 2B tested a first memory hypothesis **H1a. acquisition under intra-trial interference:** that items will be learned according to a context rule, requiring that the monkeys resolve competition among viable alternative objects in the sequence. Figures 2A and 2B present the mean learning curves and learning trial statistics, organized by screen order in the sequence, displayed for each animal separately. Both monkeys showed increased accuracy with repetition, and both learned rapidly, with a median of 4-16 trials per screen, across animals (Figure 2 AB; ^31^, see methods Behavioral analysis for details).

**Figure 2.**
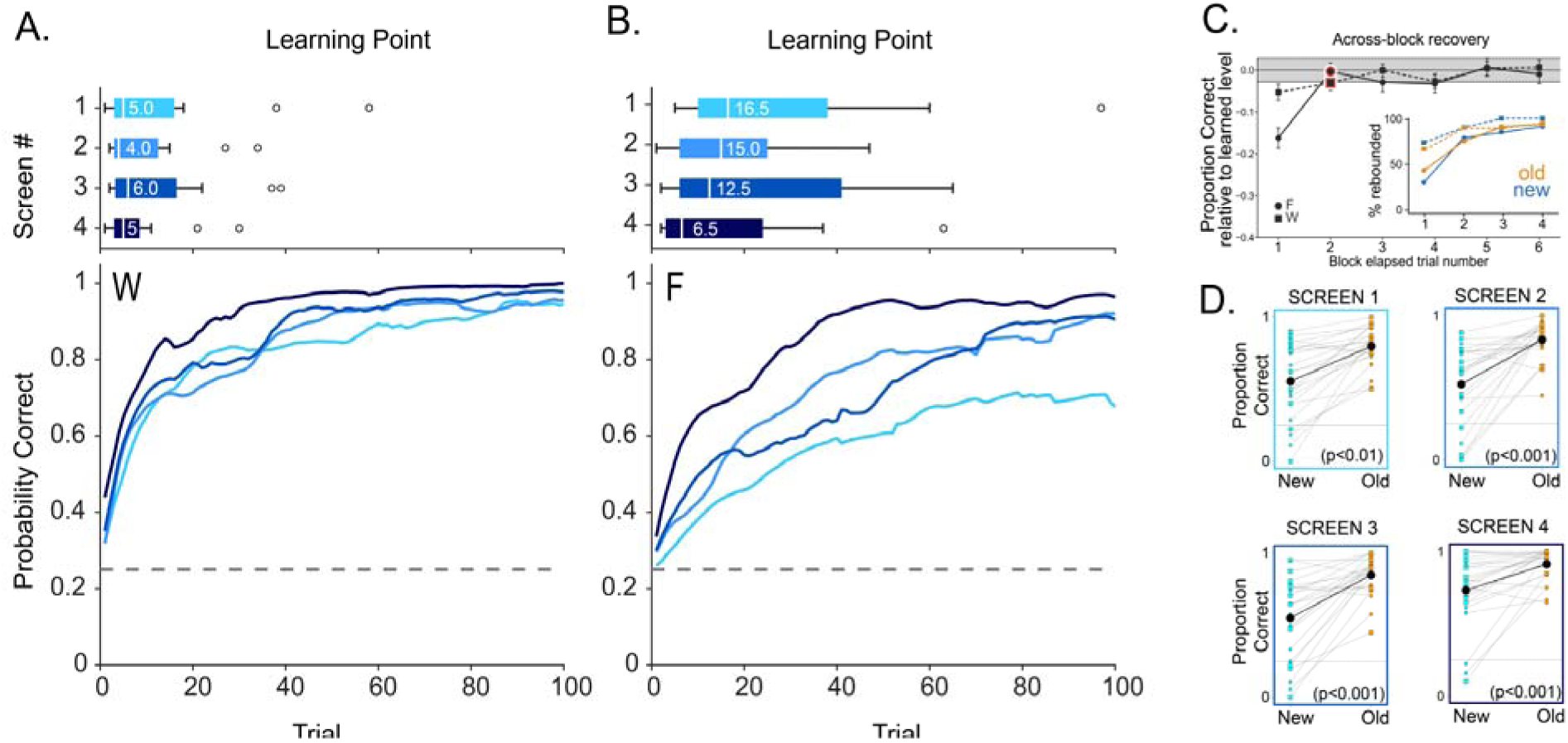
Monkeys learn sequentially presented object-context associations. (A) Top: learning points in monkey W across different sets of object-context associations, shown for each screen in the trial sequence, light-to-dark blue color-coded as in D. White bars and numbers indicate the median (N = 14 sets). Circles show outliers. Bottom: Mean learning curves across sets for each screen. The dotted grey line indicates the chance level at 0.25. (See the methods section for more details on the learning point estimation) (B) Same as A for monkey F (N = 22 sets). (C) Across-block recovery. Accuracy per trial upon returning to a set, after completing an intervening block of a different set at the other corner. Values expressed relative to the mean accuracy of the last two trials of that set’s previous block. Each point is a paired within-block difference. Error bars = SEM, N(W)=77 N(F)=67. The shaded regions are the 95% confidence interval of the mean learned level. Red icons denote the first trial no longer significantly below the learned level, sustained over the range shown. Inset: % blocks rebounded. The cumulative percentage of returning blocks back at the learned level by each of the first four trials, separated by animal and learning block type: new (blue) and old (orange).(D) Proportion of correct choices for each set on its first presentation day versus when reintroduced as an old set, shown by screen. Mean performance (black line) was higher for old than new sets on all screens (independent-samples t-test, p < 0.01 for the first screen, p<0.001 for the rest).

**H1b. Retrieval under inter-block interference:** that retrieval survives intervening blocks of competing memoranda within the session. Performance on returning to a corner sequence, as a new block begins, should rapidly resume at the level it left off at, and in fewer trials than for the initial acquisition of that material. Figure 2C shows that, for both animals, recovery to previous block levels occurred within the first 1 or 2 trials: 69% of W’s returning blocks and 38% of F’s were already recovered on the first trial; 97% and 92% by the fourth (n(W)= 77; n(F)= 67; Methods). At the group level, the first trial back fell below the level the set had reached before the interruption, by 5.3% in W (p = 0.008) and 16.2 % in F (p = 3.3e-9, one-sample t-test on within-block differences). Neither deficit outlasted that first trial back: from the second trial onward accuracy was no longer below the learned level in either animal (all later trials p >= 0.14). The same sets had taken a median of 5 and 13 trials to learn initially, so recovery after an interfering block was 5 and 6.5 times faster than acquisition (Mann-Whitney, p = 2e-22 and 2e-16).

### Memory durability

Memory under naturalistic conditions includes not only conditional selection in the moment – retrieving single experiences – but also managing the accretion of memories in the face of new experiences. This requires long-term durability, learning that is robust to proactive interference, and the ability to fluidly accommodate both learning and retrieval demands. **H1c,** memoranda will show savings even under interference from accumulated sets over weeks-long delays, evidenced through re-exposure (‘old’) performance exceeding initial acquisition, and through no drop in learning rates as sets accumulate. To assess long-term durability, monkeys performed interleaved blocks of remotely learned sets in the opposite corner from that session’s new set (see Supplement Fig. 1A). These “Old” sets had been learned several weeks prior with no intervening presentation. Performance on the old sets outpaced that of new sets, showing savings that extended to all screens, for both animals (Supplement Figure 1C; independent-samples t-test, Screen 1: t = 2.63, p < 0.01; Screen 2: t = 5.46, p < 0.001; Screen 3: t = 5.15, p < 0.001; Screen 4: t = 3.53, p < 0.001). Furthermore, on the first trial of re-exposure, accuracy over the first screens exceeded chance across sets (pooled accuracy 0.46, t(25) = 2.90, p = 0.004, one-tailed; W 0.50, t(14) = 2.56, p = 0.011; F 0.41, t(10) = 1.41, p = 0.095).

### Memory capacity

To address memory capacity, we assessed proactive interference. We tested for this by comparing learning points across the new sets presented over time. Neither animal showed worsening performance over the sets based on a linear regression of learning trial over set number, even as they demonstrated retrieval for remotely-learned material from weeks earlier. (i.e., no slowing as predicted by proactive interference): (W: t(62) = −3.85, p < 0.001; F: t(86) = 1.4; p > 0.1). On the contrary, one of the animals (W) showed significant *improvement over time,* potentially reflecting learning set or learning-to-learn.

Animals could adopt multiple approaches to learning the correct responses. These include learning four distinct spatial/temporal/visual/contextual associations, then using long-term memory to guide selection, or adopting a non-matching working memory rule that eliminates from the current choice pool any target objects from the previous screens on that trial. If animals used a non-matching working memory strategy, their likelihood of a correct choice should incrementally increase from screen 1 to 4 (Figure S1C), starting at 0.25 on screen 1 with four possible targets and reaching 1.0 on screen 4 after eliminating the three previously presented targets. Importantly, this strategy does not require long-term memory across trials, distinguishing it from associative-learning strategies that are ineffective on the first trial but become optimal with repeated exposure. To evaluate these two possibilities, we computed the proportion of success on the first trial across sets for each screen and for each animal separately (Figure S1C). Neither animal followed the rapid rise in performance across screens as expected from a working memory strategy, suggesting that long-term associative learning was the prepotent strategy.

Taken together, these results indicate that monkeys had durable, high-capacity memory that was robust to interference during challenging learning and retrieval demands.

### Head Kinematics Contain a High-Fidelity Representation of Task Variables

Having established that monkeys can learn this task while freely moving (with increased degrees of freedom), we next examined the kinematics arising from this unrestricted movement. Previous studies have shown that head movements (under full or partial head restraint) and gaze (from eye tracking) exhibit interacting neural and behavioral relationships with abstract categories and relative body position^32–34^. Building on this foundation and capitalizing on the spatial distribution of key information in the environment, we predicted that head movements would index the monkey’s place in the conditional structure, not only its place in the room, and that movement is economized where the task allows it. To determine if kinematic signals contain quantifiable information about the task, we trained a Long Short-Term Memory (LSTM) network, a type of recurrent neural network (RNN), to decode various task-related variables from the 6-axis inertial measurement unit (IMU) data during the choice epoch of the task (Figure 3A, see Methods LSTM Decoding and Dynamics for details). We trained five separate models with 10 cross validations (Figure 3B) to predict five key variables: the specific quadrant touched on the screen (one of four object choices), the screen identity as determined by its location within the enclosure (one of eight screens), the ordinal screen position within the four-step trial sequence, one of the two corners with touchscreens in the enclosure, and the trial outcome (correct or incorrect)(Figure 3C).

**Figure 3.**
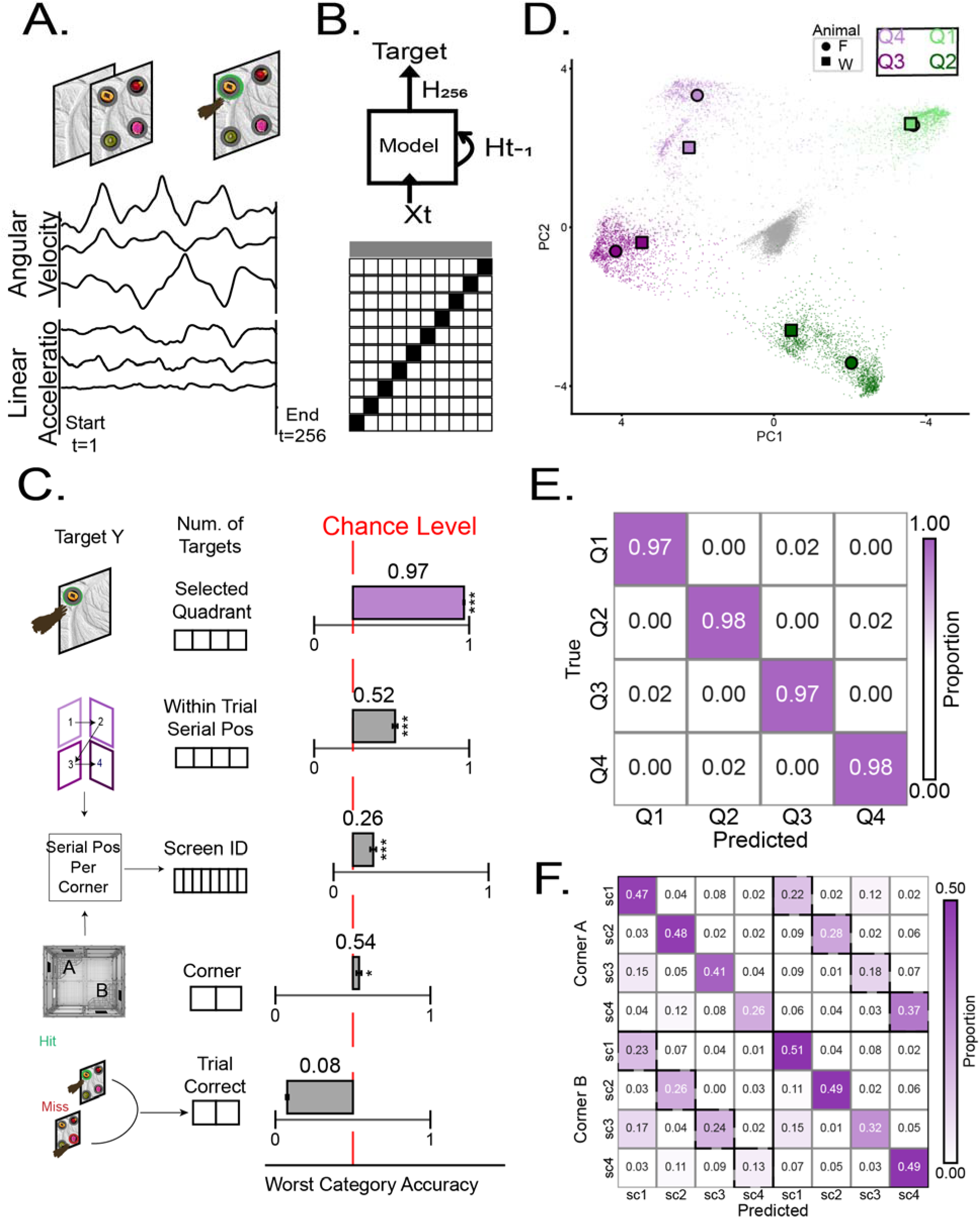
Decodable task behavior from head kinematics. (A) 6-axis IMU signal taken from the choice epoch, in which the monkeys select one of the four items on screen. The IMU time series, originally sampled at 1.5 kHz, was time-warped to the start and end of the epoch, with 256 points per epoch. (B) (***Top)*** The schematic of target variables and their corresponding model schematic; The model was a 2-layer LSTM with multiple inputs from each of the 6 IMU channels and a single output. Each target variable had its own model trained for classification. (***Bottom***) Analysis was performed in a 10-fold cross-validation, illustrated with the whole data in gray, the training fold in white, and the test fold in white (See methods for details). (C) The worst-classified category accuracy is shown for each target variable, predicted by each respective LSTM model. ***Left***: schematic of target variables for the respective model, and their number of categories in boxes. ***Right***: mean SEM worst-category accuracy across cross-validation(gray bars) aligned to chance level (dashed line, 1/number of categories) for each variable. Worst-category performance significantly exceeded chance for all variables except trial correctness (one-sample t-test, *** p < 0.001, * p < 0.05). (D) First and last time points from LSTM hidden states shown on a low-dimensional manifold via PCA with two principal components. First point is color coded in gray; last time points are color-coded by selected screen quadrant; shapes represent the quadrant centroids per animal. (E) Confusion matrix for selected quadrant position, pooled across all cross-validation test folds, and normalized by true class. Color gradients indicate high or low accuracy as shown by the color bar (F). Confusion matrix for screen ID, normalized by predicted values. The dashed off-diagonal line highlights the highest prediction errors, corresponding to screens in opposite corners at the same sequence position. Each physical corner where the screen resides is labeled on the Y-axis.

We found that head movement kinematics during the choice epoch were highly predictive of most task variables (Figure 3C). The models successfully decoded the selected quadrant within a screen, screen identity among the 8 in the trial sequence, and screen sequence position, at levels exceeding chance (one-sample t-test, p <.005). The model, however, could not reliably decode the trial outcome, and class balancing partly offsets this but still below chance level: worst-class F1 and accuracy were 0.096 and 0.796 with the natural distribution, 0.34 and 0.61 with class weighting, 0.33 and 0.6 with undersampling, and 0.34 and 0.63 with oversampling (S1 Table). Balancing lifts the worst-class score but accuracy falls to chance, indicating a variable that the representation does not index rather than an artifact of imbalance. This agrees with direct comparisons of angular velocity for error and correct trials within each session, averaging across sessions: angular velocity did not differ by outcome in either monkey (W +0.004, p = 0.18, 95% CI −0.002 to +0.010, 28 sessions; F −0.004, 0.25, 95% CI −0.011 to +0.003, 34 sessions). Head movements during the choice epoch primarily encode the animal’s intended choice, regardless of its ultimate accuracy.

### Head movements reflect abstract spatial context and sequence information

We investigated whether head kinematics also encoded broader spatiotemporal features of the task. The LSTM trained to decode the absolute physical screen identity (residing in one of eight positions) achieved a mean accuracy of 0.431±0.012, a performance significantly above the chance level of 0.125 (one-sample t-test, p <.005) (Figure 3C, F). This result indicates that a signal related to the global spatial context is present within the head movement dynamics. A detailed analysis of the decoder’s errors revealed a highly structured, non-random confusion matrix, suggesting the abstract encoding of sequence position. The model systematically confused the homotypic screens in each corner’s sequence (the Nth screen in a sequence located in one corner of the enclosure with the corresponding Nth screen of the sequence in the opposite corner: Figure 3F). This specific confusion pattern suggests that the animals employed a generic kinematic signature for each ordinal step in the sequence, which was then deployed across different spatial contexts. Unlike the decoding accuracy for the touched quadrant the decoding accuracy for screen did not increase gradually indicating early kinematics contained head movement pattern to capture spatiotemporal context of the trial (Fig S2).

Warping each epoch to a fixed number of samples removes its duration, therefore we repeated the decoding on trajectories re-referenced to real elapsed time (Fig S2). Decoding of the touched quadrant rose only in the final 300 ms before contact, crossing 50% at −500 ms, 75% at −300 ms and 90% at −200 ms in the two monkeys and reaching 0.98 and 0.97 at contact. This profile maps onto head reorientation locked to the target of executed reach rather than of a decision variable accumulating across the epoch. Screen identity also decoded, but without that late rise, and peaking at −1500ms from contact. Note that these latents come from a network trained on the *touched quadrant*, therefore that figure is a *lower bound* and is not comparable to the separately-trained screen decoder reported in Figure 3.

### Emerging gaze proxy from selected quadrant prediction

Of all the variables, the selected quadrant position was decoded with exceptionally high fidelity. The model achieved a mean accuracy of 0.975 ± 0.001 (SEM), substantially outperforming the 25% chance level (Figure 3C, E). This near-perfect performance demonstrates that a distinct and reliable kinematic signature of the head is associated with the (touched) selected quadrant position. In other words the subjects look at the item when they are making choice touch. And these high decodability was not correlated with mean angular velocity or task performance of the monkeys on the session(Supplementary Tables 3,4).

Additionally we probed the internal RNN information to understand what this model is capturing, we analyzed its internal latent space by applying Principal Component Analysis (PCA) to the final hidden state vectors from each trial. This analysis revealed that the RNN organized behavioral information into four distinct, well-separated clusters, with each cluster corresponding to one of the four screen quadrants (See Supplemental Movie, Figure 3D). The quality of this separation was quantified with a silhouette score of 0.76, indicating highly robust clustering. This representational structure was not static; the silhouette score was −0.013 at the beginning of the choice epoch, signifying choice epoch kinematics contained information for the model to predict and separate the touched quadrant. Most notably, the geometric arrangement of these clusters in the two-dimensional latent space closely resembled the physical 2×2 layout of the screen quadrants. Qualitatively (supplement video 1) and quantitatively (S4 Fig) we found that the RNN latent space corresponds to where the monkey is looking at and the latent space allows us to capture a proxy for gaze on the screen that is readily not available on a freely moving setting.

### Efficient Initial Search Pattern is Biased to the Position Nearest to the Hand

The accurate prediction of touch location and the richness of decodable kinematic features suggested gaze might be indexed even from the start of the trial. In addition, biases in visuospatial perception, attention, and decision making have been reported across species^35^; therefore, we were interested in determining whether initial head kinematics could be aligned with a gaze sweep to the first item, allowing the assessment of positional bias. All kinematic measures were derived from head movements recorded by the head-mounted IMU thus “search” here refers to gaze shifts that accompany head orienting. Quantitatively, we found that, indeed, the gaze on a specific item on the screen corresponds to the item’s cluster in latent space (See Supplemental Movie).

From the latent representations (Figure 3D) we analyzed the first 25 points (the first 10% of the warped trajectory). We fit a dynamical model to this segment (see Methods Latent Space Dynamics: for details). In real time this window spans 173 ms on the median trial for animal W and 161 ms for animal F (10% of median choice-epoch durations of 1.728 s and 1.610 s; n = 5,481 and 10,304 epochs). The same convergence is obtained when the trajectories are expressed in real elapsed time rather than in warped samples (S3 Fig). We then asked for each monkey whether we could recover the four quadrant positions, which are equally likely to be the target, or if a systematic bias would emerge in this initial sweep. In accordance with movement economy, (H3) we predicted that monkeys would choose lower-effort, shorter reach movements unless such a policy was outweighed by obtaining reward. We found that, for both animals, the dynamics of the initial movement consistently converged toward an attractor point corresponding to Quadrant 2 (Figure 4B), the bottom-right location, closest to the resting-hand positions both animals used for target selection. This demonstrates that the body matters in the shaping of the search strategy. This head kinematics serves as a proxy for overt attention, physically constrained by the body and the environment. Although not conclusive, this suggests that the resting posture creates a potential hand bias that influences the gaze path, potentially as an energy-minimization strategy.

**Figure 4.**
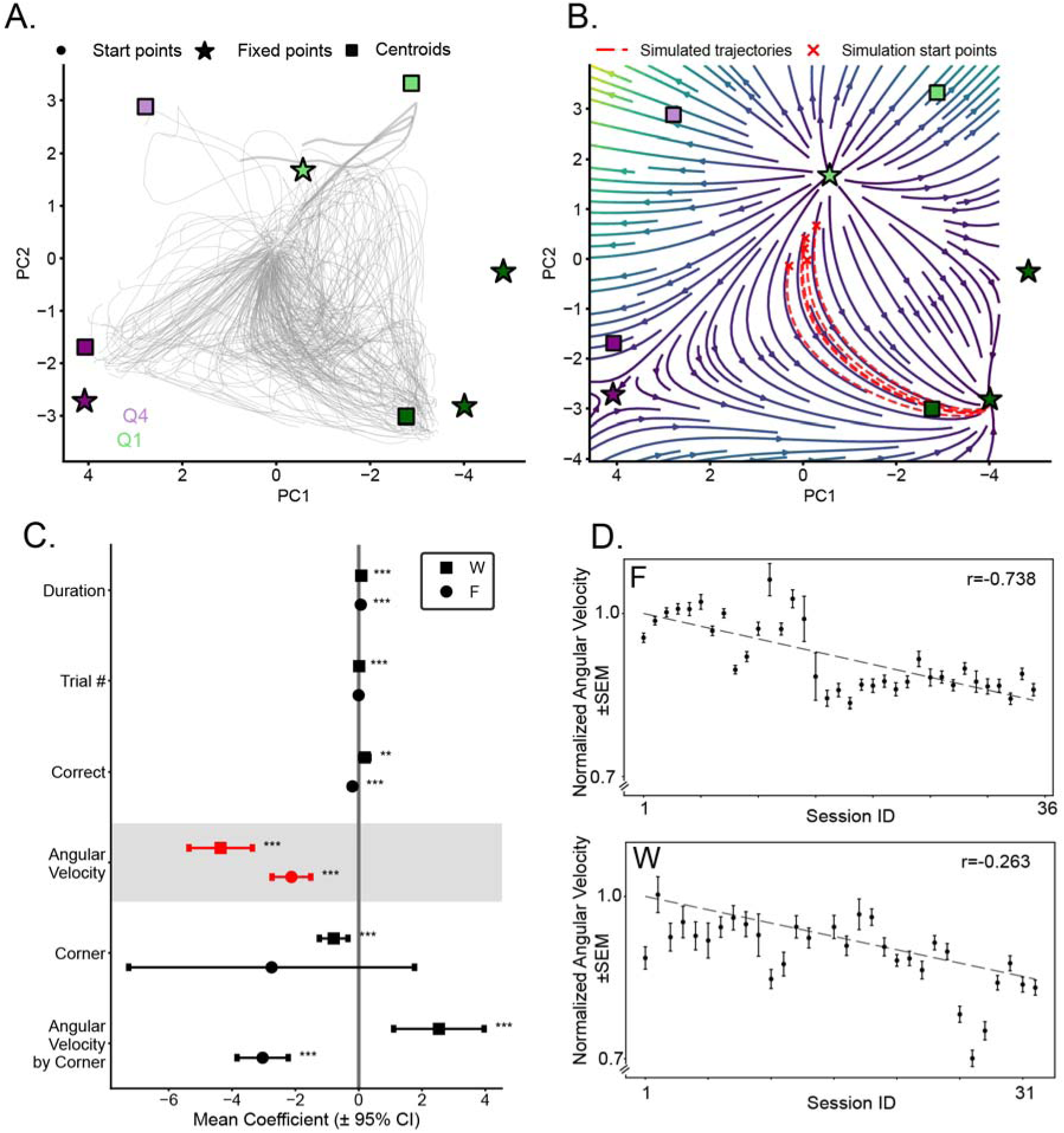
Efficient Head Kinematics. **(A)** The quadrant position LSTM model’s trajectories, early in the choice epoch, shown for the first 2 PCs of the hidden state, for Animal F. Examples of the first 25 of 256 samples of the hidden state are shown as gray lines. To characterize the geometry of these trajectories, we fit a quadratic dynamical system to each animal’s data (see Methods for details). Fixed-point attractors of the learned dynamics are shown as stars, and endpoint centroids from panel 3D are shown as squares. Both centroids and attractors are color-coded by spatial location (four choice quadrants); each attractor (star) inherits the color of its nearest trajectory endpoint centroid, determined by Euclidean distance in PC space. A total of 4 attractors were identified. **(B)** Vector field representation of the learned dynamics projected onto the first two principal components (PC1 and PC2, same projection as in A) for Animal F, evaluated on a regular grid across the PC space. The learned dynamics converge toward the attractor nearest to the negative-PC1/negative-PC2 region of the space (corresponding to the bottom-right trajectory cluster). Dashed red lines show 5 example forward simulations of 100 time steps each, initiated from empirical initial states at the start of choice epochs (red crosses). **(C)** Regression coefficients (beta weights) with 95% confidence intervals from a GLM predicting sequential session ID for each animal separately (Animals W and F; see Supplementary Tables 5, 6 for full statistics). The GLM included 8 candidate factors spanning movement, task, and performance-related variables; only factors that reached significance at p < 0.001 in at least one animal are shown here. Each animal’s coefficients are shown separately with different shapes. Angular velocity (AV) was the only factor with negative coefficient that predicted session in both animals, highlighted with red confidence intervals and gray shading. **(D)** Normalized within-session mean angular velocity ± SEM across trials within each session, plotted as a function of session index for Animals F (***Top***) and W (***Bottom***) separately. AV decreased across sessions in both, reflected in the negative regression coefficient identified in (C). SEM is computed across trials within each session.

### Head Movement Efficiency Improves Over Time

The preceding analysis reveals that monkey head movements carry a sophisticated and reliable kinematic code for task-relevant variables. Do these refined, information-rich movement patterns present from the start, or do experiences sculpt them^29,30^. We predicted that as animals master the task, their choice-related movements become more efficient overall across sessions, beyond learning the specific trial target. To test this hypothesis, we shifted our analysis from decoding information within single trials to tracking the evolution of movement kinematics across the entire learning period spanning weeks. We employed a Generalized Linear Model (GLM) to test experience-dependent changes in movement patterns. This approach allowed us to predict the ordinal session number (a proxy for accumulated experience) from kinematic parameters recorded during the choice epoch, while statistically controlling for other performance-related variables such as reaction time and trial outcome (see Methods GLM of movement dynamics). If individuals perform the task with more efficient movements across sets, independent of learning rate and accuracy, this will indicate a lower energetic cost for the same levels of task performance.

The analysis revealed a negative relationship between mean angular velocity and session number (Figure 4C), Animal W (β = −4.36) and Animal F (β = −2.12) with the effect more pronounced in Animal W, indicating that head movements became systematically slower and smoother as animals gained experience with the task. This effect remained significant even after controlling for potential confounds such as reaction time and trial outcome (correctness), confirming that the reduction in angular velocity reflects a genuine, learning-related optimization of motor strategy rather than a simple correlate of task performance. Figure 4D shows that, on average across sessions, the angular velocity decreases for both animals.

These findings demonstrate that head movement dynamics undergo systematic changes with task experience during the choice epoch, reflecting an increase in the efficiency of actions during deliberation with expertise.

## DISCUSSION

The present study demonstrates that freely moving macaques can acquire and retain numerous conditional object–in–context sequences in a multi-level 3-D enclosure. Across dozens of four-step sequences, each object served as the rewarded target in only one specific spatiotemporal position and was a distractor elsewhere, yet under these conditions both animals learned 56-88 unique item-context associations. Retrieval also proved robust to competing material interleaved during learning. On returning to a set within the same session, across intervening blocks of another set, both animals rapidly returned to their prior levels of performance, Typically on the first or second trial. Interference within the session therefore produced a brief cost rather than relearning. Furthermore, they showed robust savings when remotely learned sets were reintroduced after weeks-long delays, while continuing to acquire more memoranda without evidence of proactive interference. This extends lesion ^8,36–38^ and neurophysiological evidence^12,14,39^ for relational object-in-context and scene memory in primate medial temporal networks in an ecologically richer, sequence-structured environment.

Platforms for unrestrained primate behavior have made a range of natural activity measurable, including navigation, foraging, action repertoires and social interaction^55,57,59,60,62,63^. Macaques have been tasked with learning according to previous reward locations in an arena^5,6^ foraging patches^62^ and from a social partner^59^. Our task differs from these in the kind of contingency it imposes. The animal must retrieve which object belongs at a given step of a four-screen sequence, at eight stations whose local demands are identical, having accumulated 50 to 80 such associations on eight reused contexts. The judgment is renewed at every screen, and it cannot be resolved by position on the screen or by the value of a ‘patch’.

Our tests were inspired by sequence-of-events frameworks developed in rats, posing the same conditional question, in settings where movements are restricted in order to sample stimulus cues^40^.

The task design also incorporates elements of immersive memory tasks in humans, in which items are embedded in a structured spatial route or environment and retrieved by traversing that route/imagining the spatial configuration. Enhanced recall is generally, though not always, associated with such immersive, actively explored surroundings ^18,19^. Here, monkeys physically traverse a stable sequence of screens distributed across levels and corners of the enclosure, selectively binding the objects as targets specified by their spatiotemporal positions rather than merely recognizing them in isolation. This aligns with evidence that hippocampal, prefrontal and posteriormedial systems encode temporal context and event sequences^40–46^, generalize across memories sharing item–context structure^47,48^ and support scene-based relational memory ^49–52^, while situating these computations in a setting closer to everyday primate behavior than traditional immobile tasks using a single screen display with 2-D visuospatial arrays.

A key motivation for the Treehouse environment was to reconcile naturalistic behavior with the demands of high-density wireless primate electrophysiology ^53, 56^. Many spatial or relational tasks that move toward ecological validity, such as open-field navigation, provide limited repetitions of any given condition, restricting the ability to align neural activity to repeated, precisely timed decision events, and limiting trial-or event-averaged analyses. In contrast, the present task combines rich spatial context and extended routes with sufficient repetition and tight temporal control: stimuli appear on fixed touchscreens with precisely timed cue and choice epochs, while animals move freely between vertically and horizontally separated stations that approximate natural foraging and exploration. This structure is compatible with wireless neural recordings, building on work from this group and others describing in freely-moving and sleeping macaques hippocampal sharp-wave ripples and other state-dependent oscillations^54–56^, sleep reactivation^56^, gaze or body spatial modulation ^39,57^, and decision and movement variables ^58–63^. Where behaviors and neural recordings are compatible across species, this additionally allows comparisons with rodent neurophysiology conducted in similarly immersive or navigational settings. The treehouse task and screen location are fully reconfigurable, opening up future possibilities for asking different cognitive questions across different kinematic settings.

While performing the task, the monkeys expressed structured and increasingly efficient head-movement patterns that tightly link cognition, spatial context, and motor control. By instrumenting the head with a lightweight IMU, we leveraged the spatial separation of screens, and of objects on the screens, to treat head kinematics as a continuous readout of the location the animal selects. An LSTM trained solely on 6-axis IMU signals during the choice epoch decoded the selected screen quadrant with near-perfect accuracy. We probed into the “black-box” latent geometry and found that it recapitulated the physical 2×2 layout of the on-screen array *<u>without any alignment across animals</u>*, despite differing sensor orientations across animals. Beyond the selected quadrant, the same kinematic signals reflected additional variables apart from reach. Holding ordinal step and touched quadrant constant, the screen position was still decodable (balanced accuracy 0.613 and 0.621 against a shuffled control of 0.506 and 0.491), and within a single ordinal step the screen was still recoverable (0.625 to 0.573 across steps in one animal, 0.709 to 0.556 in the other). Ordinal step, in turn, generalized across the enclosure: a classifier trained at one corner read the step at the other with sessions held out at 0.472 and 0.466 against 0.505 and 0.515 within corner, retaining 84% of within-corner performance where the label-shuffled null is 0.251. The screen decoder’s errors are consistent with this, systematically confusing homotypic steps across corners, but the two variables are separable. These results converge with prior work on eye–head–hand coordination and active vision in head-unrestrained primates ^21,66–68^, where coordinated movements optimize visual sampling and future reaches, and they demonstrate that IMU-based decoding can provide a practical, implant-compatible alternative or complement to head-mounted eye-tracking systems ^68^. The readout was limited in this experiment: trial outcome did not decode from the IMU signal, even with class balancing. The signals report the location the animal selects, and where it is in the task or in the space, rather than whether the selection was correct.

A central contribution of this work is to evaluate movement organization that may optimize more functions than just the task-required movements, including sensitivity to the neuroeconomics of effort and energy. The initial search dynamics inferred from latent LSTM geometry provide a complementary window onto effort-sensitive gaze policies without calibrated eye tracking that is not feasible under freely moving setting. At the beginning of the choice epoch, head-movement trajectories for both monkeys converged reliably toward the quadrant closest to the resting hand, rather than sampling all positions evenly. This consistent bias toward the least costly reach direction indicates a policy adopted under uncertainty rather than a perceptual bias: because target position is balanced across the four locations, no positional preference can raise the reward rate, so reaching to the nearest position neither costs nor earns accuracy and is abandoned only as retrieval becomes reliable. Movement kinematics may reveal latent valuations of effort and benefit that are otherwise difficult to access^21,64^. Our findings extend this logic to whole-head movements in a memory task: both the early bias toward near-hand quadrants and the across-session decline in angular velocity indicate that head kinematics may be adjusted according to the state of retrieval and the cost of reaching, with no measured contribution from expected reward.

Across sessions, mean head angular velocity during the choice epoch decreased systematically, irrespective of accuracy, indicating a shift toward efficient head movement as animals gained experience with the overall task mechanics. This decline follows session order rather than exposure to a set, and does not restart when new sets are introduced, so it reflects growing familiarity with the enclosure rather than progress on any particular material. This experience-dependent reduction in movement speed without loss of accuracy is consistent with the idea that motor strategies are optimized to reduce energetic cost as internal models of the task improve seen in other motor-skill settings. The movement-vigor literature has framed similar phenomena in terms of an internal trade-off between expected reward and the energetic or subjective cost of movement: saccade vigor and limb-movement speed scale with the subjective value of options and with the history of reward and effort^27,28^, and metabolic measures can be used to quantify how physical effort devalues prospective rewards^69,70^. In our task, the fact that angular velocity declines with experience, while performance remains high, suggests that as the conditional rules become familiar, the subjective benefit of additional, high-vigor movements diminishes relative to their energetic cost, pushing the system toward more economical policies.

More broadly, these results support the view that primate memory for objects, contexts, and sequences could in principle build embodied contents, that may emerge from the interplay between hippocampo-cortical representations and species-typical patterns of active sensing^71–74^. The Treehouse environment provides a scalable, electrophysiology-ready structure for which whole-body and head kinematics may furnish a continuous, energetically tuned readout of choice formation, sequence context, and movement economy. This platform will enable future experiments to ask how hippocampal–cortical ensembles represent conditional item–in–context rules as the memory space grows, and more generally, how neural circuits track learning-to-learn and decision-linked movement vigor. Together, the present findings show that monkeys can express capacious, interference-robust sequence memory while freely moving through a controlled yet naturalistic environment, and they situate the kinematics of the head as a high-fidelity signature of cognitive state governed by the same cost–benefit principles that link vigor, effort, and decision making in humans ^75–78^.

## Supporting information

Video of head movement latents

## ACKNOWLEDGMENTS

The authors want to thank the veterinary staff for maintaining the well-being of animals, Daicia Allen, Mitchell Riley, and Chrissy Suell for animal training support, and Sergey Motorny, Supeng Wu, and Richard Song for their technical support for pose tracking.

## Author contributions

KLH and SA conceptualized the study. KLH and WZ designed the apparatus. WZ, KLH, and SA developed the experimental design. KLH and SA collected the data. SA and KFR curated the data. SA, KFR, HT, and KLH conducted the formal analysis. SA, KLH, and KFR prepared visualizations for data presentation. SA KFR and KLH wrote the original draft. WZ contributed to the review and editing. KLH provided funding for the study and supervised the project. All authors discussed and interpreted the results.

## Funding

This work was funded by the Whitehall Foundation, NEI P30EY008126, and NI R01 NS127128.

## DECLARATION OF INTERESTS

Conflict of interest. The authors declare that the research was conducted in the absence of any commercial or financial relationships that could be construed as a potential conflict of interest.

## DECLARATION OF GENERATIVE AI AND AI-ASSISTED TECHNOLOGIES

During the preparation of this work, the author used Cursor to accelerate analysis, enhance visualization, and improve the efficiency of the analysis pipeline. After using this tool or service, the author(s) reviewed and edited the content as needed and take full responsibility for the content of the publication.

## MATERIALS AND METHODS

All animal procedures were approved by the Vanderbilt Institutional Animal Care and Use Committee, in compliance with the policies of the United States Department of Agriculture and Public Health Service on the humane care and use of laboratory animals. Experimental subjects were two adult female rhesus macaque monkeys (*Macaca mulatta*).

### Behavioral Testing

Testing environment. The testing apparatus consisted of a custom-made enclosure (1.52 x 1.52 x 2.13 m; the ‘Treehouse’, Figure 1a, b), equipped with modular panels organized into upper and lower levels. Among the panels, 8 were designated as testing stations, each equipped with a touchscreen (ViewSonic 24” 1080p 10-Point Multi Touch Screen Monitor, models 2421 and 2455) and a perch. These stations were arranged symmetrically in two opposed corners (e.g., northwest and southeast) with a 2-level x 2-side arrangement in each corner. Fluid reward was delivered by peristaltic pumps (Campden Instruments Precision Liquid Feed Pump) placed in both active corners. The pumps were controlled with a Measurement Computing Corporation USB DAQ (OM-USB-1208FS) that was connected to the Experimental Control System.

Behavioral tracking. We monitored and recorded animals’ behavior in using 8 side cameras and 1 overhead camera (Figure 1a; 5 e3Vision, WhiteMatter cameras, https://white-matter.com/products/e3vision/, and 4 Logitech webcams) set up around the enclosure. We note that the camera array was positioned for whole-enclosure coverage rather than per-screen detail, with the animal facing away from or orthogonal to cameras, so scored gaze was not available, and reach trajectories were incomplete due to occlusion. Video frames from all cameras/webcams were collected at 30Hz with HD resolution. e3Vision frames from all cameras were sent to the e3Vision hub and synchronized online before recording. Logitech video frames were synced to e3Vision frames manually by aligning a reference frame (a simultaneous change of background on all screens). In addition to video recordings, we recorded movements, with a wireless 9-axis Inertial Measurement Unit (IMU, Freelynx, Neuralynx, Inc.) to previously implanted titanium cranial fixtures (Double-Asymmetric Head Post, Gray Matter Research or similar custom fixtures from Rogue Research). The IMU data were acquired at 3kHz and saved directly to the acquisition computer’s disk. To synchronize IMU and camera frames, TTL pulses for the start and end of the e3Vision frames were sent from the e3Vision hub to the acquisition system where IMU data were recorded.

### Experimental control

Experiments were controlled by a single computer equipped with two AMD graphics cards (Cape Verde PRO FirePro W600) that connected to the eight touchscreens, plus one control monitor. A MATLAB toolbox based on PLDAPS ^79^ was developed in house (‘TreeTop’ at https://github.com/hoffman-lab/TreeHouse)^80^. This toolbox uses Psychtoolbox functionality ^81,82^) to control stimulus selection and presentation on each monitor independently with precise timing, of stimulus presentations and monitoring multi-touch activations of the screens with exact locations, and control of reward delivery. To achieve the precision and independence in stimulus timing and touch monitoring, the toolbox relies on object oriented programming using a finite state system approach (^80^) as outlined for the PLDAPS ^79^ and Opticka packages ^83^. The TreeTop system also sent event codes to the IMU DAQ, allowing experimental control, cameras, and IMU to be synchronized.

### Trial structure

As shown in Figure 1C and D, the task started with the presentation of a start cue (black circle) that designated the active screen for the animal. Upon touching the cue, and after a 1 s delay, the array onset consisted of the appearance of 4 objects presented in a 2 by 2 grid on a background scene providing context specific for that set of objects. Selection: If monkeys touched the correct item (“Target”) - the item associated with that screen - a ‘correct’ tone was played, and the target reappeared at the center of the screen. Confirmation: The subject had to touch the target again to proceed to activate the next screen. If, instead, she selected one of the 3 non-target objects (“distractors”), an ‘error’ tone was played along with the disappearance of the stimuli for 2-4 seconds. Then, the correct target was shown in isolation in the originally presented location in the background context (i.e., Correction). Monkeys had to touch the target in isolation to proceed to the confirmation where they had to select the target once again. Both animals selected targets with the right hand; a frame-by-frame check of one full session of each monkey confirmed that every task touchscreen touch was right-handed. Occasionally, only the target was displayed, with no distractors, to help maintain motivation. These were equally likely across screens and were excluded from all learning analyses. As shown in Figure 1D, subjects proceeded through the trial sequence on all 4 screens, in order, before receiving fluid reward. For Subject W, the total reward was calculated based on the performance of the animal within a trial (e.g., 2 drops of juice per correct selection and 0 drops for incorrect touches) and delivered at the reward receptacle. For subject F, a fixed reward amount was delivered, but the reward type (flavor) was switched in the middle of the session to maintain motivation. For the presented data, both subjects always traversed an identical sequence starting with screen 1 (upper left) to screen 4 (lower right) on one corner of the environment.

### Session design

Although a given stimulus set was restricted to one corner of the apparatus, sets could be assigned to either of the two touchscreen corners. Within each session the two corners alternated in blocks (S1 Fig), which allows retrieval to be measured both at the first encounter of the day and on returns to a corner following a block of an intervening set. For monkey W, only one corner contained a new set; the other corner contained a set that had been learned previously. Presentation of new sets was staggered across corners in this way. For monkey F, earlier sessions consisted of two new sets on opposite sides of the apparatus, but later sessions were similar in setting to monkey W. We continued training on the same sets across days until monkeys learned the novel sets and then introduced new different sets.

### Visual stimuli

Objects and background images were chosen from colored photorealistic fruits/vegetables and natural scenes, respectively. The isolated object images were scaled to approximately the same size and placed on a black circle background of 73 mm diameter with a gray ring of 117 mm diameter around it, to make objects distinct from the screen background image (Figure 1). Different types of fruits/vegetables were included in any a set of four objects, and images were never reused across sets.

### Experimental subject pretraining

First, the monkeys were pretrained to use touch screens. Subject W was then required to select by touching a correct synthetic object paired with a specific scene background in the booth setup, while seated in a transport chair. Subject F was required to touch one of two possible item colors, or one of two spatial positions as a function of the screen location screen in the treehouse. When training started in the Treehouse, both monkeys were therefore naïve to screen-contingent object association tasks but were familiar with learning selection rules on touchscreens. After the animals learned the location of all the reward spouts, we introduced an operant cue touch on a single screen for reward, followed by selection from among an increasing number of items in an array. Finally, we increased the number of active screens in which the animal had to perform the same actions, but across the multiple screens in a predetermined order before receiving reward until a sequence of all 4 screens from a given corner was achieved.

### Behavioral analysis

Learning assessment. We concatenated all completed trials per stimulus set, including all responses in which the animals selected from among the 4 objects in the array. We then used a latent-process model to calculate individual learning curves for each screen of each set, by subject ^31^. The process includes a 2-step state-space filtering followed by a smoothing algorithm to estimate the learning curve and its confidence intervals. The estimation used an Expectation Maximization algorithm with an initial background probability of 0.25 and convergence criterion of 1e-8. The maximum number of steps was set to 5000, and a significant epsilon (square root of the variance of the learning state process) of 0.005 was used. The initial conditions were fixed, and the learning trial was defined for each animal as the first trial where the 95% confidence interval of the estimated probability of correct performance exceeded and remained above chance. Three sets were excluded based on poor model convergence. To estimate changes in learning rate across sets (i.e., over time) for each animal, we fit a linear regression to the learning rate of each screen of each set, according to the ordinal position of the set.

Across-block recovery. Blocks of consecutive trials alternated corners and corresponding stimulus sets (S1 Fig). We evaluated returns: visits to a set the animal had already worked earlier the same day, with a block of the other set in between. We used as a reference level the mean accuracy over the last two trials of that set’s previous visit, measured before the interruption. Returns were excluded if the previous visit was shorter than six trials or where the reference level did not exceed chance by 0.2, on the grounds that regaining a level that was never learned is not retrieval (2 and 3 returns in F; 0 and 2 in W). Each trial of the returning block was compared with its own block’s reference level by a one-sample t-test on the within-block differences. Recovery is reported at a group-level criterion: the first trial at which the mean difference across blocks was no longer below the reference level, and remained so over the six trials analyzed. We also report recovery as a per-block count: the median trials to regain the level, indicating the typical number of trials a single return took to reach its specific reference. New and old material were compared on both reported measures (Mann-Whitney) and pooled. Blocks were truncated at twelve trials and the first six are shown. Recovery is compared against initial acquisition using the learning trial value, so the two thresholds differ in the conservative direction: the learning trial is the first trial whose confidence interval clears chance, a lower bar than the left-off level a return has to regain.

### IMU tracking

The IMU data tracked the linear acceleration and angular velocity of head movement. Angular velocity measures roll, pitch, and yaw, and linear acceleration measures linear movements. These data can augment behavioral assessments by providing higher temporal resolution (3000 Hz). Our interest was in the relative movement of the head at decision making part of trial. We extracted IMU data, in microvolts, from the choice epoch of each trial, (I.e., the cue touch to selection). During this time, the animals need to choose among four options, and this choice is the target of learning in the present experiments. Each IMU axis was scaled by its per-axis maximum, computed over the whole session. Whole sessions were held out of training, therefore this scaling is within-session and cannot transfer information between folds. A classifier trained on odd sessions and tested on even sessions predicted the selected quadrant at 0.981 across 7,139 trials (0.988 in F, 0.973 in W; S1 Table).

### LSTM Decoding

To quantify the information content of head kinematics, a variant RNN called LSTM was employed to decode task-related variables from the 6-axis IMU data (3 axes of linear acceleration, 3 axes of angular velocity). We used an LSTM model because we were working with time series and were interested in exploring the temporal evolution of head movement, along with decoding. All analyses focused on the “choice epoch,” defined as the interval from the appearance of the choice array to the animal’s touch selection. For each trial, the IMU time series within this epoch was time-normalized to a fixed length of 256 samples. To test whether this normalization discarded information carried by duration, the same latent trajectories were re-referenced to real elapsed time. A logistic classifier was then fitted at each time point on the ten leading latent principal components, with whole sessions held out (five-fold, grouped on session). Trials were restricted to a reaction time of at least 2 s so that cohort composition was fixed across time points (n = 1,365, median 2.48 s in F; n = 971, 2.68 s in W), and accuracy was evaluated on a 100 ms grid..

The full dataset comprised 15,785 trials from both animals. We trained separate models to predict distinct categorical task variables. Model performance was evaluated using a 10-fold cross-validation procedure, and we report the mean accuracy across the test folds. The network architecture consisted of a standard Long Short-Term Memory (LSTM) model with two layers and 128 hidden units. The model was trained to predict a single categorical output using the Adam optimizer with a learning rate of 0.001.

To investigate the network’s learned representations, we performed a latent space analysis on the model trained to predict the selected quadrant. For this analysis, the model was trained on data from even-numbered sessions (8,646 trials) and validated on data from odd-numbered sessions (7,139 trials). The final hidden state vector from each validation trial was extracted, and its dimensionality was reduced for visualization using Principal Component Analysis (PCA). To quantify the separability of the choice-related clusters, silhouette scores were calculated on the first 3 principal component of the hidden states at both the first and final time points of the epoch.

### Latent Space Dynamics

For the dynamic analysis in Figure.4, we began by extracting the first 25 samples—10% of the 256-sample choice epoch—from every trial and analysed each monkey separately. These time-series segments were projected onto the first two principal components to obtain a compact state vector x(t). We then uncovered the governing head-movement dynamics with the Sparse Identification of Non-linear Dynamics (SINDy)^85^ algorithm, using a second-order polynomial dictionary and sequential thresholded least-squares regression to obtain a parsimonious set of differential equations x = f(x)Evaluating f(x) on a dense grid across the empirical PC space yielded vector-field and streamline visualizations, while fixed points were located by numerically solving f(x) = 0 from multiple random seeds. Each attractor was assigned a behavioral label by mapping it to the nearest centroid of the task-dependent hidden-state clusters defined earlier. Finally, we seeded the learned dynamical system with every trial’s empirical initial state. We integrated the equations for 25 time steps, recording the attractor and trajectory for each simulation. This procedure revealed how early-epoch head kinematics rapidly converge onto a small set of stable, behaviorally meaningful attractors as animals master the task.

### Generalized Linear Model (GLM) of Movement Dynamics

To assess how movement patterns changed as a function of long-term experience, we analyzed trial-level statistics using a Generalized Linear Model (GLM). For each choice epoch, we calculated the root mean square (RMS) of the angular velocity and linear acceleration, along with other task-related variables (Table 1).

**Table 1.** Task variable predictors for GLM\.

| Variable | Explanation |
|---|---|
| Angular Velocity RMS | The root mean square of the three angular velocity components: sqrt(AVx<sup>2</sup> + AVy<sup>2</sup> + AVz<sup>2</sup>) |
| Linear Acceleration RMS | The root mean square of the three linear acceleration components:sqrt(LAx<sup>2</sup> + LAy<sup>2</sup> + LAz<sup>2</sup>) |
| Correct | Binary variable indicating whether the choice was correct (1) or incorrect (0) |
| Trial Number | The ordinal trial number within the current session |
| Duration | The reaction time for the choice epoch, in seconds |
| Corner Bin | Categorical variable indicating whether the trial occurred in a novel or familiar corner |

All analyses were conducted in Python using the PyTorch, Scikit-learn, SciPy, and NumPy libraries. Visualizations were generated using Matplotlib and Plotly.

We constructed two separate ordinal GLMs, one for each animal, with the session number as the dependent variable. The model’s predictors included the kinematic and task variables listed below. This approach enabled us to isolate the effect of long-term learning (session number) on movement dynamics while controlling for variations in trial-to-trial performance. For visualization, the mean and SEM of the Angular Velocity RMS were calculated for each session and normalized for each animal to highlight the rate of decrease across learning.

---

**Supplementary Video 1.** Dynamics of gaze: projection from LSTM latents. Examples of monkey during the choice epoch, alongside the concomitant principal component projection of the latents from the LSTM model of head movements.

**Supplementary Figure 1**: (A) Schematic illustrating how stimulus sets transition from New to Old status across experimental sessions. (B) Distribution of the number of days between when a set is last presented as New and when it reappears as Old labeled as memory lag (N=20). (C) The proportion of correct choices on the first trial across sets for each screen for both animals (monkey W = square and monkey F = circle); the dashed line shows the chance level at 0.25. The dotted grey line with square markers indicates the expected performance if monkeys were using only a non-match, working memory strategy across items in the trial sequence, such that at each progressing sequence element, there would be one less item of choice

**Supplementary Figure 2:**Decoding of touched quadrant (left) and screen identity (right) from the head signal in real (non-warped) time. Latent trajectories were re-referenced from the warped 256-sample epoch to real elapsed time, and a logistic classifier was fitted at each time point on the 10 leading latent PCs, with sessions held out (5-fold, grouped on session number). Balanced accuracy; chance = 1/k (dotted horizontal line). Dashed vertical lines mark the first time durably above chance (gray) and, where a ceiling exists, the crossings of 0.50, 0.75 and 0.90.

**Supplementary Figure 3:** Warped latent space dynamic vs actual time latent space dynamics (**A**) and real time 0–0.16 s with dt = 10 ms (**B**); Fig 4’s window spans 0.172 s on this animal’s median trial.

**Supplementary Figure 4:** The latent trajectory visits options other than the one selected, animals combined. The three panels of the previous figure with both animals’ 7,139 trials pooled into a single sample; every measure and convention is the same. Each count in the steps per trial reflects the centroid a trajectory’s position is best aligned with. Chance is 25% or 63.75 steps. “Selected” means the option the animal touched. A) Steps per trial directed at the selected option’s centroid, at the Q2 centroid, and at the remaining two centroids; *steps toward Q2 are counted only on trials where Q2 was not itself the choice, so the three categories do not overlap*. B) Distribution across trials of the proportion of steps directed at the selected option; solid line, the pooled mean (33.9%); dashed line, chance. As expected the selected option draws a weighted share of the gaze, when collapsing time/warp steps elapsed in the trial. C) Time-resolved shares. The proportion rises across the epoch that ends with the monkey’s selection (touch). Pooled, only 33.9% of steps point at the option eventually selected — above the 25% chance level, but far from the near-100% reach implied, with about 66% pointing at one of the three alternatives. Tertiles: 27.4% to 31.9% to 42.4%.

**S1 Table — selected-quadrant decoding across a session boundary**

**S2 Table — class balancing and trial-outcome decodability**

**S3 Table – decoding vs mean AV**

**S4 Table – decoding vs behavioral accuracy**

**S5 Table:** GLM Summary for Monkey F

**S6 Table:** GLM Summary for Monkey W

## Notes

### Competing Interest Statement

The authors have declared no competing interest.

### Summary of Updates

This revision added head kinematic analysis and additional memory assessments.

## REFERENCES

1. Bachevalier, J., Nemanic, S. & Alvarado, M. C. The influence of context on recognition memory in monkeys: effects of hippocampal, parahippocampal and perirhinal lesions. Behav. Brain Res. 285, 89–98 (2015).

2. Browning, P. G. F. & Gaffan, D. Global retrograde amnesia but selective anterograde amnesia after frontal-temporal disconnection in monkeys. Neuropsychologia 46, 2494–2502 (2008).

3. Froudist-Walsh, S. et al. The Rhesus Monkey Hippocampus Critically Contributes to Scene Memory Retrieval, But Not New Learning. J. Neurosci. 38, 7800–7808 (2018).

4. Gaffan, D. Scene-specific memory for objects: a model of episodic memory impairment in monkeys with fornix transection. J. Cogn. Neurosci. 6, 305–320 (1994).

5. Hampton, R. R., Hampstead, B. M. & Murray, E. A. Selective hippocampal damage in rhesus monkeys impairs spatial memory in an open-field test. Hippocampus 14, 808–818 (2004).

6. Lavenex, P. B., Amaral, D. G. & Lavenex, P. Hippocampal lesion prevents spatial relational learning in adult macaque monkeys. J. Neurosci. 26, 4546–4558 (2006).

7. Murray, E. A. & Mishkin, M. Object recognition and location memory in monkeys with excitotoxic lesions of the amygdala and hippocampus. J. Neurosci. Off. J. Soc. Neurosci. 18, 6568–6582 (1998).

8. Parkinson, J. K., Murray, E. A. & Mishkin, M. A selective mnemonic role for the hippocampus in monkeys: memory for the location of objects. J. Neurosci. 8, 4159–4167 (1988).

9. Zola, S. M. et al. Impaired recognition memory in monkeys after damage limited to the hippocampal region. J. Neurosci. Off. J. Soc. Neurosci. 20, 451–463 (2000).

10. Erickson, C. A. & Desimone, R. Responses of macaque perirhinal neurons during and after visual stimulus association learning. J. Neurosci. 19, 10404–10416 (1999).

11. Gulli, R. A. et al. Context-dependent representations of objects and space in the primate hippocampus during virtual navigation. Nat. Neurosci. 23, 103–112 (2020).

12. Hussin, A. T., Abbaspoor, S. & Hoffman, K. L. Retrosplenial and Hippocampal Synchrony during Retrieval of Old Memories in Macaques. J. Neurosci. 42, 7947–7956 (2022).

13. Jutras, M. J., Fries, P. & Buffalo, E. A. Gamma-band synchronization in the macaque hippocampus and memory formation. J. Neurosci. 29, 12521–12531 (2009).

14. Montefusco-Siegmund, R., Leonard, T. K. & Hoffman, K. L. Hippocampal gamma-band Synchrony and pupillary responses index memory during visual search. Hippocampus 27, 425–434 (2017).

15. Naya, Y. & Suzuki, W. A. Integrating what and when across the primate medial temporal lobe. Science 333, 773–776 (2011).

16. Robertson, R. G., Rolls, E. T. & Georges-Fran ois, P. Spatial view cells in the primate hippocampus: effects of removal of view details. J. Neurophysiol. 79, 1145–1156 (1998).

17. Wirth, S. et al. Single neurons in the monkey hippocampus and learning of new associations. Science 300, 1578–1581 (2003).

18. Maidenbaum, S. et al. Improved spatial memory for physical versus virtual navigation. J. Neural Eng. 22, 046014 (2025).

19. Smith, S. A. Virtual reality in episodic memory research: A review. Psychon. Bull. Rev. 26, 1213–1237 (2019).

20. van Helvoort, D., Stobbe, E., Benning, R., Otgaar, H. & van de Ven, V. Physical exploration of a virtual reality environment: Effects on spatiotemporal associative recognition of episodic memory. Mem. Cognit. 48, 691–703 (2020).

21. Diamond, J. S., Wolpert, D. M. & Flanagan, J. R. Rapid target foraging with reach or gaze: The hand looks further ahead than the eye. PLoS Comput. Biol. 13, e1005504 (2017).

22. Song, J.-H. & Nakayama, K. Hidden cognitive states revealed in choice reaching tasks. Trends Cogn. Sci. 13, 360–366 (2009).

23. Draschkow, D., Kallmayer, M. & Nobre, A. C. When Natural Behavior Engages Working Memory. Curr. Biol. 31, 869–874.e5 (2021).

24. Findlay, J. M. & Gilchrist, I. D. Active Vision: The Psychology of Looking and Seeing. xiii, 220 (Oxford University Press, New York, NY, US, 2003). doi:10.1093/acprof:oso/9780198524793.001.0001.

25. Krajbich, I., Armel, C. & Rangel, A. Visual fixations and the computation and comparison of value in simple choice. Nat. Neurosci. 13, 1292–1298 (2010).

26. Adolph, K. E. & Avolio, A. M. Walking infants adapt locomotion to changing body dimensions. J. Exp. Psychol. Hum. Percept. Perform. 26, 1148–1166 (2000).

27. Nikooyan, A. A. & Ahmed, A. A. Reward feedback accelerates motor learning. J. Neurophysiol. 113, 633–646 (2015).

28. Summerside, E. M., Shadmehr, R. & Ahmed, A. A. Vigor of reaching movements: reward discounts the cost of effort. J. Neurophysiol. 119, 2347–2357 (2018).

29. Yadav, G. & Duque, J. Reflecting on what is ‘skill’ in human motor skill learning. Front. Hum. Neurosci. 17, 1117889 (2023).

30. Marin Vargas, A., et al. Task-driven neural network models predict neural dynamics of proprioception. Cell 187, 1745–1761.e19 (2024).

31. Smith, A. C. et al. Dynamic analysis of learning in behavioral experiments. J. Neurosci. 24, 447–461 (2004).

32. C Pallus, A. & G Freedman, E. Target position relative to the head is essential for predicting head movement during head-free gaze pursuit. Exp. Brain Res. 234, 2107–2121 (2016).

33. Mok, R. M. & Love, B. C. Abstract Neural Representations of Category Membership beyond Information Coding Stimulus or Response. J. Cogn. Neurosci. 34, 1719–1735 (2022).

34. Saeb, S., Weber, C. & Triesch, J. Learning the optimal control of coordinated eye and head movements. PLoS Comput. Biol. 7, e1002253 (2011).

35. Odegaard, B., Wozny, D. R. & Shams, L. Biases in Visual, Auditory, and Audiovisual Perception of Space. PLOS Comput. Biol. 11, e1004649 (2015).

36. Gaffan, D. Normal forgetting, impaired acquisition in memory for complex naturalistic scenes by fornix-transected monkeys. Neuropsychologia 31, 403–406 (1993).

37. Murray, E. A., Baxter, M. G. & Gaffan, D. Monkeys with rhinal cortex damage or neurotoxic hippocampal lesions are impaired on spatial scene learning and object reversals. Behav. Neurosci. 112, 1291–1303 (1998).

38. Templer, V. L. & Hampton, R. R. Cognitive mechanisms of memory for order in rhesus monkeys (Macaca mulatta). Hippocampus 23, 193–201 (2013).

39. Xu, X., Du, K. & Mao, D. Spatial dissociation between recognition and navigation in the primate hippocampus. Sci. Adv. 10, eado7392 (2024).

40. Allen, T. A., Morris, A. M., Mattfeld, A. T., Stark, C. E. L. & Fortin, N. J. A Sequence of events model of episodic memory shows parallels in rats and humans. Hippocampus 24, 1178–1188 (2014).

41. Eichenbaum, H. Time cells in the hippocampus: a new dimension for mapping memories. Nat. Rev. Neurosci. 15, 732–744 (2014).

42. Fortin, N. J., Agster, K. L. & Eichenbaum, H. B. Critical role of the hippocampus in memory for sequences of events. Nat. Neurosci. 5, 458–462 (2002).

43. Hsieh, L.-T., Gruber, M. J., Jenkins, L. J. & Ranganath, C. Hippocampal activity patterns carry information about objects in temporal context. Neuron 81, 1165–1178 (2014).

44. Manns, J. R., Howard, M. W. & Eichenbaum, H. Gradual changes in hippocampal activity support remembering the order of events. Neuron 56, 530–540 (2007).

45. Shima, K., Isoda, M., Mushiake, H. & Tanji, J. Categorization of behavioural sequences in the prefrontal cortex. Nature 445, 315–318 (2007).

46. Xie, Y. et al. Geometry of sequence working memory in macaque prefrontal cortex. Science 375, 632–639 (2022).

47. Jang, Y. & Huber, D. E. Context retrieval and context change in free recall: recalling from long-term memory drives list isolation. J. Exp. Psychol. Learn. Mem. Cogn. 34, 112–127 (2008).

48. Long, N. M. & Kahana, M. J. Successful memory formation is driven by contextual encoding in the core memory network. Neuroimage 119, 332–337 (2015).

49. Chrastil, E. R. Heterogeneity in human retrosplenial cortex: A review of function and connectivity. Behav. Neurosci. 132, 317–338 (2018).

50. Kaplan, R. et al. Medial Prefrontal-Medial Temporal Theta Phase Coupling in Dynamic Spatial Imagery. J. Cogn. Neurosci. 29, 507–519 (2017).

51. Maguire, E. A., Intraub, H. & Mullally, S. L. Scenes, spaces, and memory traces: what does the hippocampus do? The Neuroscientist 22, 432–439 (2016).

52. Ranganath, C. & Ritchey, M. Two cortical systems for memory-guided behaviour. Nat. Rev. Neurosci. 13, 713–726 (2012).

53. Parodi, F., Kording, K. P. & Platt, M. L. Primate neuroethology: a new synthesis. Trends Cogn. Sci. 0, (2025).

54. Abbaspoor, S., Hussin, A. & Hoffman, K. L. Theta- and gamma-band oscillatory uncoupling in the macaque hippocampus. BioRxiv 10.1101/2021.12.30.474585 (2022) doi:10.1101/2021.12.30.474585.

55. Talakoub, O., et al. Hippocampal and neocortical oscillations are tuned to behavioral state in freely-behaving macaques. BioRxiv 10.1101/552877 (2019) doi:10.1101/552877.

56. Abbaspoor, S., Aljishi, A. & Hoffman, K.L. Experience reorganizes content-specific memory traces in macaques. Nat Neurosci 29, 1976–1986 (2026). 10.1038/s41593-026-02357-2

57. Mao, D. et al. Spatial modulation of hippocampal activity in freely moving macaques. Neuron 109, 3521–3534.e6 (2021).

58. Berger, M., Agha, N. S. & Gail, A. Wireless recording from unrestrained monkeys reveals motor goal encoding beyond immediate reach in frontoparietal cortex. eLife 9, e51322 (2020).

59. Franch, M. et al. Visuo-frontal interactions during social learning in freely moving macaques. Nature 627, 174–181 (2024).

60. Lanzarini, F. et al. Neuroethology of natural actions in freely moving monkeys. Science 387, 214–220 (2025).

61. Lu, H.-Y. et al. Distinct tasks engage a shared neural subspace in human hippocampus and anterior cingulate cortex. 2026.04.24.720703 Preprint at 10.64898/2026.04.24.720703 (2026).

62. Shahidi, N. et al. Population coding of strategic variables during foraging in freely moving macaques. Nat. Neurosci. 27, 772–781 (2024).

63. Testard, C., Tremblay, S. & Platt, M. From the field to the lab and back: neuroethology of primate social behavior. Curr. Opin. Neurobiol. 68, 76–83 (2021).

64. Korbisch, C. C., Apuan, D. R., Shadmehr, R. & Ahmed, A. A. Saccade vigor reflects the rise of decision variables during deliberation. Curr. Biol. 32, 5374–5381.e4 (2022).

65. Marshall, T. R., Ruesseler, M., Hunt, L. T. & O’Reilly, J. X. The representation of priors and decisions in the human parietal cortex. PLOS Biol. 22, e3002383 (2024).

66. Arora, H. K. et al. Eye-head-hand coordination during visually guided reaches in head-unrestrained macaques. J. Neurophysiol. 122, 1946–1961 (2019).

67. Freedman, E. G. & Sparks, D. L. Eye-head coordination during head-unrestrained gaze shifts in rhesus monkeys. J. Neurophysiol. 77, 2328–2348 (1997).

68. Singh, V. P., Li, J., Dawson, K., Mitchell, J. F. & Miller, C. T. Active vision in freely moving marmosets using head-mounted eye tracking. Proc. Natl. Acad. Sci. U. S. A. 122, e2412954122 (2025).

69. Bruening, G. W., Courter, R. J., Sukumar, S., O’Brien, M. K. & Ahmed, A. A. Disentangling the effects of metabolic cost and accuracy on movement speed. PLoS Comput. Biol. 20, e1012169 (2024).

70. Huang, H. J., Kram, R. & Ahmed, A. A. Reduction of Metabolic Cost during Motor Learning of Arm Reaching Dynamics. J. Neurosci. 32, 2182–2190 (2012).

71. Martinez-Trujillo, J. Why do primates have view cells instead of place cells? Trends Cogn. Sci. 29, 226–229 (2025).

72. Meister, M. L. R. & Buffalo, E. A. Getting directions from the hippocampus: The neural connection between looking and memory. Neurobiol. Learn. Mem. 134, 135–144 (2016).

73. Murray, E. A., Wise, S. P. & Graham, K. S. Representational specializations of the hippocampus in phylogenetic perspective. Neurosci. Lett. 680, 4–12 (2018).

74. Rolls, E. T. & Wirth, S. Spatial representations in the primate hippocampus, and their functions in memory and navigation. Prog. Neurobiol. 171, 90–113 (2018).

75. Phillips, M. H. & Edelman, J. A. The dependence of visual scanning performance on saccade, fixation, and perceptual metrics. Vision Res. 48, 926–936 (2008).

76. Movement vigor as a traitlike attribute of individuality. J. Neurophysiol. 120, 741–757 (2018).

77. Shadmehr, R., Huang, H. J. & Ahmed, A. A. A Representation of Effort in Decision-Making and Motor Control. Curr. Biol. 26, 1929–1934 (2016).

78. Sim, M. & Kim, J.-U. Differences between experts and novices in kinematics and accuracy of golf putting. Hum. Mov. Sci. 29, 932–946 (2010).

79. Eastman, K. M. & Huk, A. C. PLDAPS: A hardware architecture and software toolbox for neurophysiology requiring complex visual stimuli and online behavioral control. Front. Neuroinformatics 6, 1 (2012).

80. Wagner, F. Modeling Software with Finite State Machines: A Practical Approach. (Auerbach Publications, 2006). doi:10.1201/9781420013641.

81. Brainard, D. H. The Psychophysics Toolbox. Spat. Vis. 10, 433–436 (1997).

82. Pelli, D. G. The VideoToolbox software for visual psychophysics: transforming numbers into movies. Spat. Vis. 10, 437–442 (1997).

83. Andolina, I. M. Opticka: Psychophysics-toolbox based experiment manager. Zenodo 10.5281/zenodo.7787740 (2023) doi:10.5281/zenodo.7787740.

84. Sang-Ah Yoo, R. Shayna Rosenbaum, John K. Tsotsos, Mazyar Fallah, Kari L. Hoffman; Long-term memory and hippocampal function support predictive gaze control during goal-directed search. Journal of Vision 20(5):10. 10.1167/jov.20.5.10 (2020).

85. S.L. Brunton, J.L. Proctor, & J.N. Kutz,Discovering governing equations from data by sparse identification of nonlinear dynamical systems, Proc. Natl. Acad. Sci. U.S.A. 113 (15) 3932–3937, 10.1073/pnas.1517384113 (2016).

